# JAGGER localization and function is dependent on GPI anchor addition

**DOI:** 10.1101/2023.11.06.565777

**Authors:** Raquel Figueiredo, Mónica Costa, Diana Moreira, Miguel Moreira, Jennifer Noble, Luís Gustavo Pereira, Paula Melo, Ravishankar Palanivelu, Sílvia Coimbra, Ana Marta Pereira

## Abstract

In flowering plants, successful double fertilization requires the correct delivery of two sperm cells to the female gametophyte inside the ovule. The delivery of a single pair of sperm cells is achieved by the entrance of a single pollen tube into one female gametophyte. To prevent polyspermy, Arabidopsis ovules avoid the attraction of multiples pollen tubes to one ovule – polytubey block. In Arabidopsis *jagger* mutants, a significant number of ovules attract more than one pollen tube to an ovule due to an impairment in synergid degeneration. *JAGGER* encodes a putative arabinogalactan protein (AGP) which is predicted to be anchored to the plasma membrane by a glycosylphosphatidylinositol (GPI) anchor. Here, we show that JAGGER fused to citrine yellow fluorescent protein (JAGGER-cYFP) is functional and localizes mostly to the periphery of ovule integuments and transmitting tract cells. We further investigated the importance of GPI-anchor addition domains for JAGGER localization and function. Different JAGGER proteins with deletions in predicted ω-site regions and GAS (GPI attachment signal) domain, expected to compromise the addition of the GPI anchor, led to disruption of JAGGER localization in the cell periphery. All JAGGER proteins with disrupted localization were also not able to rescue the polytubey phenotype, pointing to the importance of GPI-anchor addition to *in vivo* function of the JAGGER protein.

## INTRODUCTION

Plant reproduction involves a series of cell-cell interactions between the male gametophyte and the female gametophyte and between the male gametophyte and the female sporophyte (Johnson et al. 2019; Lopes et al. 2019). In Arabidopsis, the embryo sac (female gametophyte) is surrounded by the ovule maternal tissues, and in addition to the two female gametes (the egg cell and the central cell), it contains five accessory cells: two synergid cells and three antipodal cells (Yadegari and Drews 2004). After the pollen lands on the stigma, it produces a tube and carries the two sperm cells at its tip, and grows past the stigma, style, transmitting tract, before emerging in the ovary. The tube continues to grow in the ovary and when it is attracted to the ovule, it gains access to the embryo sac through one of the synergid cells, which facilitates the double fertilization of the egg cell and the central cell by each of the two sperm cells carried by the pollen tube, giving rise to the embryo and the seed endosperm, respectively (Russell 1992).

A functional female gametophyte is essential for the attraction of the pollen tube to the ovule (Shimizu and Okada 2000; Johnson et al. 2019; Lopes et al. 2019). In *in vitro* experiments with *Torenia fournieri*, in which the micropylar end of the female gametophyte protrudes outside the ovule integuments, the pollen tubes were attracted to LURE-type CRP attractant peptides from the synergids (Higashiyama et al., 1998; Takeuchi, 2021; Zhong et al., 2019). The two synergid cells located at the micropylar end of the ovule share a specialized secretory structure called the filiform apparatus, which is rich in cell wall, plasma membrane, and secretory vesicles (Huang and Russell 1992). The highly invaginated filiform apparatus vastly increases the plasma membrane surface area that could come in quick and easy contact with the arriving pollen tube and promotes the secretion of small peptides and glycosides involved in the pollen tube attraction, reception, and subsequent release of the sperm cells for double fertilization (Li et al. 2018; Pereira and Coimbra 2019).

The first pollen tube receptor discovered in the synergid filiform apparatus was the receptor-like protein kinase FERONIA/SIRENE (FER/SRN). Unlike in wild type, the *fer*/*srn* mutation in ovule results in incessant growth of the pollen tube within the embryo sac, and consequently male gametes end up not being released from the pollen tube (Escobar-Restrepo et al. 2007). LORELEI (LRE) is a putative glycosylphosphatidylinositol (GPI) anchored protein (GPI-AP) that is primarily expressed in the synergids and the *lre* mutant ovule phenocopy all the aberrant behaviors of *fer*/*srn* mutant ovule (Capron et al. 2008). LRE and FER are believed to be co-receptors and work together at the interface between pollen tube and synergid cells to regulate the rupture of the pollen tube in the synergid cells (Liu et al. 2016). Arabinogalactan proteins (AGPs) are highly glycosylated proteins and are likely involved in several steps of the reproductive process. The AGPs were found to be attractants and/or signalling molecules for pollen tube guidance in *Nicotiana tabacum*, as it displayed a gradient of increasing glycosylation in the extracellular matrix of the transmitting tract, from the stigma to the style, and enabled pollen tube growth and guidance (Cheung et al. 1995; Wu et al. 1995). As evidenced by immunolocalization studies in ovules of different species, AGPs are present in the sporophytic integument cells and in the synergid cells (Costa et al., 2013; Coimbra et al., 2007; Coimbra and Salema, 1997; Lopes et al., 2016; Pereira et al., 2016b, 2016a; Hou et al., 2016; Lara-Mondragón and MacAlister, 2021). Remarkably, in *T. fournieri*, an AGP derived sugar (AMOR) produced by the ovule sporophytic tissues induced competency of the pollen tube to respond to the LURE-type CRP attractant peptides from the synergids (Mizukami et al. 2016). It has been suggested that given the importance of Ca^2+^ dynamics for pollen tube growth and the ability of AGPs to bind Ca^2+^ in a pH-dependent manner, AGPs might influence pollen tube growth either by acting as a periplasmic Ca^2+^ reservoir or as a diffusible Ca^2+^ carrier (Lamport et al. 2021). In the ovary of Arabidopsis, a group of chimeric AGPs known as early nodulin-like proteins (ENODLs 11-15) are highly expressed in the embryo sac and their loss of function disrupts the bursting of the pollen tubes upon penetration (Hou et al. 2016). The influence of AGPs on pollen tube growth has also been observed in several species using the Yariv reagent that precipitates AGPs (Qin et al. 2007; Lara-Mondragón and MacAlister 2021). Another important AGP is AGP4 or JAGGER, which plays an important role in preventing polytubey (Pereira et al. 2016b). The failure to prevent polytubey in the *jagger* mutant could be due to the continued secretion of pollen tube attractants from the synergid cell that persists after fertilization instead of being rapidly inactivated. Indeed, the persistent synergid in *jagger* mutant remains aberrantly active after fertilization and does not undergo degeneration, resulting in enhanced frequency of polytubey (Pereira et al. 2016b).

Approximately half of the AGP family members in Arabidopsis, including JAGGER, are predicted to contain a GPI lipid anchor that directs the proteins to the outer leaflet of the plasma membrane, making them worthy candidates for signal perception and transduction (Seifert and Roberts 2007; Ellis et al. 2010). The functional properties of AGPs probably reside in their variable sugar chains (Moreira et al. 2023). These sugar chains may become easily accessible as signalling molecules through detachment of the AGPs from the cell wall and its secretion to the apoplast by the action of phospholipases (GPI-PLC) (Majewska-Sawka and Nothnagel 2000; Ellis et al. 2010). In eukaryotes, the newly synthesized proprotein chain of GPI-APs typically contain an N-terminal signal peptide (SP) for translocation into the endoplasmic reticulum (ER) during its synthesis, and an extended C-terminal GPI-attachment signal (GAS) region containing several discernible elements: a region of about 11 amino acid residues plus two segments that are critical for addition of the GPI anchor: i) a cleavable hydrophobic C-terminal tail of 9-24 amino acid residues and ii) a ω-site region containing four small residues, one of which will be attached to a GPI anchor. The ω-site region and the C-terminal tail are separated by a spacer region of about 6 amino acid residues (Eisenhaber et al. 2003).

To date, the structure has been resolved for only one plant GPI anchor, and it is that of *Pyrus communis* AGP1 (PcAGP1). It was shown to contain a glycan moiety, composed of three α-linked mannose residues and one glucosamine that is linked to the C-terminus of the protein by ethanolamine phosphate, and a lipid moiety of phosphoceramide, composed of phytosphingosine and tetracosanoic acid and attached to the glycan core by myo-inositol (Oxley and Bacic 1999; Yeats et al. 2018; Desnoyer and Palanivelu 2020). The GPI anchor is synthesized in the ER and is then attached to the ω-site of GPI-APs by the GPI transamidase complex. After GPI anchor attachment to the ω-site of GPI-APs, the glycan and lipid moieties are remodeled. After leaving the ER, GPI-APs are transported by the secretory pathway, through the Golgi complex, to their final destination, the plasma membrane (Ellis et al. 2010; Yeats et al. 2018). Disrupting GPI anchor synthesis or attachment in Arabidopsis is often gametophytic or embryo lethal, as assessed by loss-of-function mutant studies of genes that code for specific subunits of the GPI transamidase complex (Bundy et al. 2016; Desnoyer et al. 2020). Indeed, many GPI-APs have important functions in plant reproduction, such as AGP4 (JAGGER), AGP6, AGP11, and AGP18, FLA3, ENODL 11-15 and LRE that are important for Arabidopsis gametophyte development or fertility (Acosta-García and Vielle-Calzada, 2004; Coimbra et al., 2009; Li et al., 2010; Hou et al., 2016; Liu et al., 2016; Pereira et al., 2016a). Other examples include LORELEI-LIKE GPI-anchored proteins 2 and 3, which are involved in maintaining cell wall integrity at the pollen tube tip during its growth through the style and transmitting tract (Ge et al. 2019; Feng et al. 2019), and COBRA-LIKE protein 10 (COBL10), which seems to be important for micropylar guidance of the pollen tube to the synergids (Li et al. 2013).

Despite these critical roles of specific GPI-APs in both gametophytes, the importance of subcellular localization of GPI-APs in the plasma membrane via a GPI anchor is not fully understood and remains to be shown that they are essential for their functions. Indeed, while mutations in the LRE GAS region or the loss of *trans*-acting GPI transamidase resulted in mutant LRE proteins or wild type LRE protein, respectively, not localized in the plasma membrane-rich filiform apparatus of synergid cells, they however did not affect LRE protein function in pollen tube reception (Liu et al. 2016). These surprising results showing that GPI anchoring is dispensable for LRE function in the female gametophyte could not still be reconciled with the evolutionary conservation of domains required for GPI anchor attachment in LRE proteins. So, in the present work we investigated whether loss of GPI anchoring is dispensable in JAGGER, another GPI-AP. For this, we generated JAGGER protein with several deletions that are expected to compromise the addiction of the GPI anchor, verified if JAGGER subcellular localization is disrupted in cells of ovules, and examined if these changes reversed the *jagger* mutant phenotype. Our work, besides describing the subcellular localization of an AGP *in planta*, demonstrated that GPI anchor is critical for *in vivo* function of JAGGER.

## MATERIAL AND METHODS

### JAGGER sequence analysis

The *JAGGER*-*cYFP* DNA constructs were designed as described for LRE in (Liu et al. 2016). JAGGER ω-sites were predicted by the BIG-PI plant predictor software (https://mendel.imp.ac.at/gpi/plant_server.html) (Eisenhaber et al. 2003).

### Plant Material

*Arabidopsis thaliana* seeds were surface sterilized and plated on half strength Murashige and Skoog medium, containing 2% (w/v) sucrose and the appropriate antibiotics. After stratification at 4°C for 2-3 days, seeds were placed in a growth chamber and maintained at 20°C in a 16 h/8 h photoperiod. Seven-to ten-days-old seedlings were transferred to soil and kept in the same growth conditions. The background ecotype of all Arabidopsis seeds was Columbia (Col-0). The *jagger1-2* (N412778) mutant line was previously characterized by (Pereira et al. 2016b).

### JAGGER - cYFP DNA constructs

The *cYFP* (E1403; Tian et al., 2004) reporter gene was fused to wild-type *JAGGER* in the C-terminus and expressed in transgenic Arabidopsis plants under the control of *JAGGER* promoter (2273 bp). cYFP was used as reporter because it is pH insensitive and capable of fluorescing in the acidic apoplast (Gjetting et al. 2012), which is the likely site of localization of GPI-APs (Schultz et al. 1998). cYFP has been used to determine plasma membrane localization of other Arabidopsis GPI-anchored surface proteins (Simpson et al. 2009; Liu et al. 2016). The inclusion of a complete GAS sequence downstream of the cYFP sequence is expected to make the fusion protein GPI-anchored. Flanking the cYFP reporter, two equal JAGGER ω-11 sequences were inserted. Upstream of the cYFP is the native ω-11 sequence and the second ω-11 domain was placed between the cYFP and the ω-site to prevent any potential hinderance to the GPI anchoring mechanism and the need for regulatory cues. Care was taken to retain the same amino acid sequence of the ω-11 sequence in JAGGER, yet the codons were altered to be consistent with the codon bias in Arabidopsis. The JAGGER-cYFP construct was structured in this manner, as cleaving the AGP for fusion in the ω-11 keeps the coding sequence unaltered and should not potentially impair the JAGGER and cYFP protein folding.

Three DNA fragments were prepared by PCR amplification as follows: 1) a genomic sequence encompassing *JAGGER* regulatory region and CDS up to the codon immediately upstream of the predicted ω-site; 2) *cYFP* sequence preceded by a Gly^9^ linker sequence (Chen et al. 2013) corresponding to GRPGGGGGA, which was amplified from plasmid DNA; 3) a codon-optimised ω-11 sequence followed by the predicted ω-site and the 3’-terminal sequence corresponding to the C-terminal end region of JAGGER. The primers were designed to allow the fusion of the three fragments by overlap PCR (Table S1). The *JAGGER-cYFP* fusion under the control of the native promoter was cloned into a modified pH7WG (Karimi et al. 2002; Liu et al. 2016). This vector has inherited an unique *AscI* restriction site from *pENTR/D-TOPO* that was used together with *SpeI* restriction site to linearize the vector and clone the *proJAGGER:JAGGER-cYFP* construct using In-Fusion HD Cloning Plus (Clontech). For the mutant constructs used in this study, altered-sequence fragments were introduced by PCR using modified primers (Table S1) and *AscI/SpeI* linearized *pH7WG::ProJAGGER:JAGGER-cYFP* was used as template for in-fusion cloning.

### Isolation of insertion lines

*pH7WG::ProJAGGER:JAGGER-cYFP* and the *ProJAGGER:JAGGER-cYFP* mutant constructs plasmids were transferred to *Agrobacterium tumefaciens* (GV3101 strain) and plants (wild-type and *jagger1-2* mutant line) were then transformed by the floral dip procedure (Clough and Bent 1998). Hygromycin-resistant transformants were selected and verified by PCR with primers flanking the *JAGGER-cYFP* coding sequence (primers described in Table S1) followed by Sanger sequencing of the resultant PCR products.

### Confocal imaging

Fluorescent images were captured using a Leica TCS SP8 confocal laser scanning microscope (CLSM). For cYFP imaging, samples were excited with a 514 nm laser line, and emission spectra between 522 and 550 nm were collected. cYFP images were processed with ImageJ software (http://imagej.nih.gov/ij/). Pistils from flowers at stage 12 according to (Smyth et al. 1990) were dissected under a stereomicroscope (Model C-DSD230, Nikon) using hypodermic needles (0.4 x 20 mm; Braun). The opened carpels and the ovules that remained attached to the septum were kept in water and covered with a coverslip.

### Preparation of live plant material for microscopy

For crosses with dehiscent anthers, closed flower buds were emasculated 24–48 h before pollination, at stage 11–12, according to (Smyth et al. 1990). Fresh pollen at flower stage 13 was profusely applied to the stigma of the emasculated pistils. Hand-pollinated pistils were collected 2 days after pollination and used for aniline blue staining.

### Aniline blue pollen tube staining

Arabidopsis pistils were fixed overnight in 10% (v/v) acetic acid in ethanol at 4°C, washed three times with water and softened with 8 M NaOH for 16 h, washed three times with water, and incubated overnight in 0.1% (w/v) decolorized aniline blue (DAB) at 4°C (Kho and Baër 1968). Stained pistils were dissected under a stereomicroscope (Model C-DSD230, Nikon) using hypodermic needles (0.4 × 20 mm; Braun). The opened carpels and the ovules that remained attached to the septum were maintained in a drop of DAB and covered with a coverslip. Observations were made with a Leica DMLB/LB30T at 350-400 nm (UV), and polytubey was scored on a Nikon DS-Ri2 camera using the NIS-Elements BR program (version 4.60.00).

### Statistical Analysis

Statistical tests on the data were performed using GraphPad software (www.graphpad.com).

## RESULTS

### JAGGER-cYFP localizes in the cell periphery of transmitting tissue and ovule integument cells in Arabidopsis pistils

The JAGGER amino acid sequence contains putative features that define a GPI-AP: the N-terminal signal peptide (SP), the proline-rich yet a less complex ω-11 region, and two domains that are key to the addition of a GPI anchor: the ω-site region and the C-terminal hydrophobic tail (GAS sequence) (Figure 1). The putative GAS sequence in JAGGER has the conserved features of animal GAS sequences (Schultz et al. 1998) and is predicted to span from residue Ala^118^ through to Leu^134^. JAGGER Ser^111^ was the best predicted ω-site residue by BIG-PI plant predictor software (Eisenhaber et al. 2003).

**Fig. 1.**
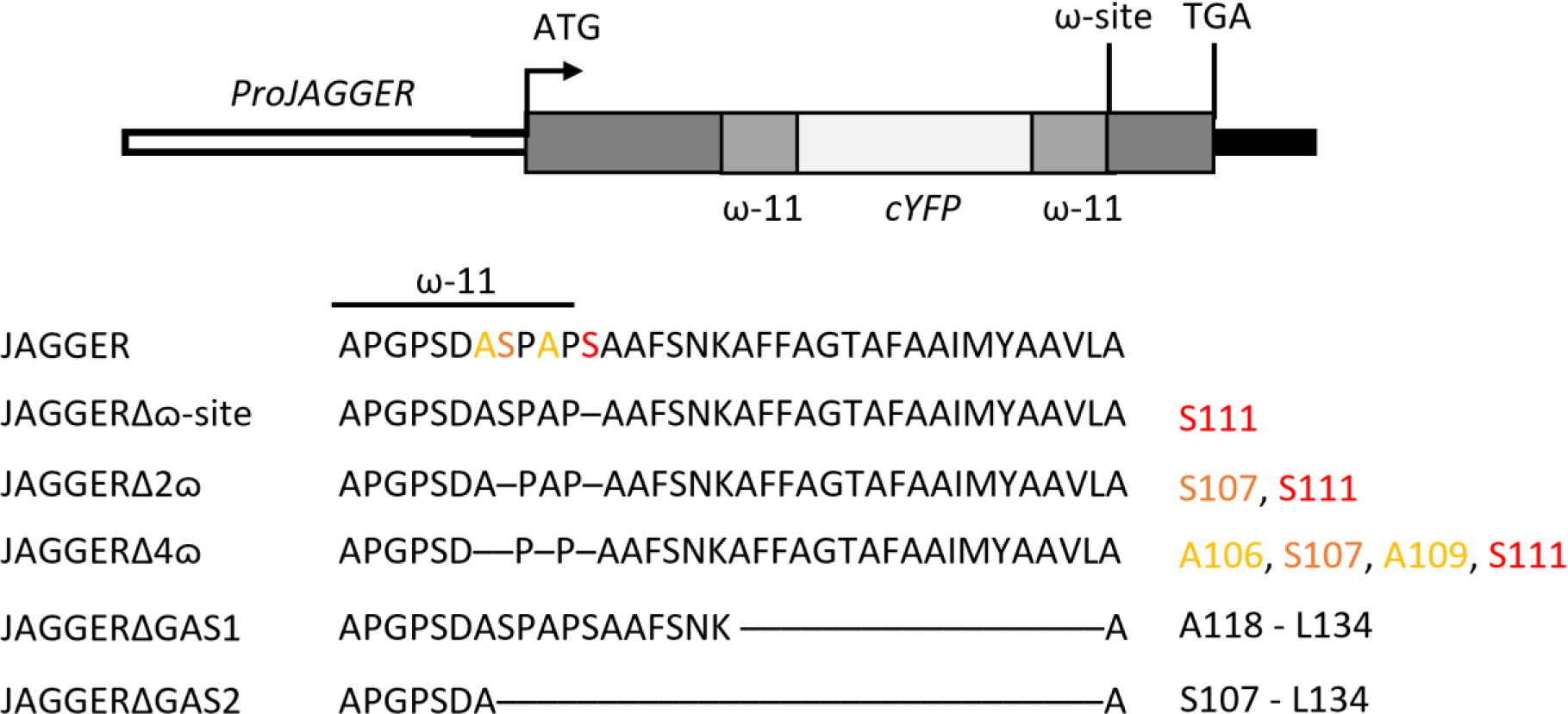
Diagram of the JAGGER-cYFP protein Wild-type or altered amino acid sequence on either side of the predicted ω-site amino acid are indicated below the diagram. In JAGGER, the best predicted ω-site and a cryptic ω-site are labelled in red and orange, respectively. Two additional cryptic ω-sites are labelled yellow. Each dash in the protein sequence represents a deletion of the corresponding amino acid in wild-type JAGGER protein sequence. The corresponding amino acid is referred at right. (Diagram not drawn to scale.)

To study subcellular localization of JAGGER in the Arabidopsis pistil, and to test if the translational fusion constructs can complement the *jagger* mutant phenotype, we fused the citrine YFP (cYFP) reporter gene to the C-terminus of JAGGER coding sequence and expressed it from the *JAGGER* promoter. JAGGER protein and cYFP amino acid sequences were separated by a semi-flexible linker of 9 amino acids, adapted from the one used in LORELEI-cYFP (Liu et al. 2016) and included in all DNA constructs described in this work.

Confocal laser scanning microscopy (CLSM) images of unpollinated pistils carrying the wild-type JAGGER protein fused to cYFP showed that JAGGER-cYFP fusion proteins were expressed in the sporophytic ovule integument cells and in the transmitting tissue cells (Figure 2). Ovule integument cells located more closely to the embryo sac region presented a higher level of fluorescence intensity compared to the chalaza region of the ovules (Figure 2 A – B). Also in these cells, some cYFP fluorescence is detected in puncta in the cell’s cytoplasm. Isolated transmitting tract cells revealed cYFP signal mostly at the cell periphery (Figure 2 C – D) and did not overlap with the auto-fluorescent regions of the cell that likely represented the chloroplasts. In both cell types, cYFP signal was detected at the cell periphery, consistent with JAGGER being a putative GPI-AP.

**Fig. 2.**
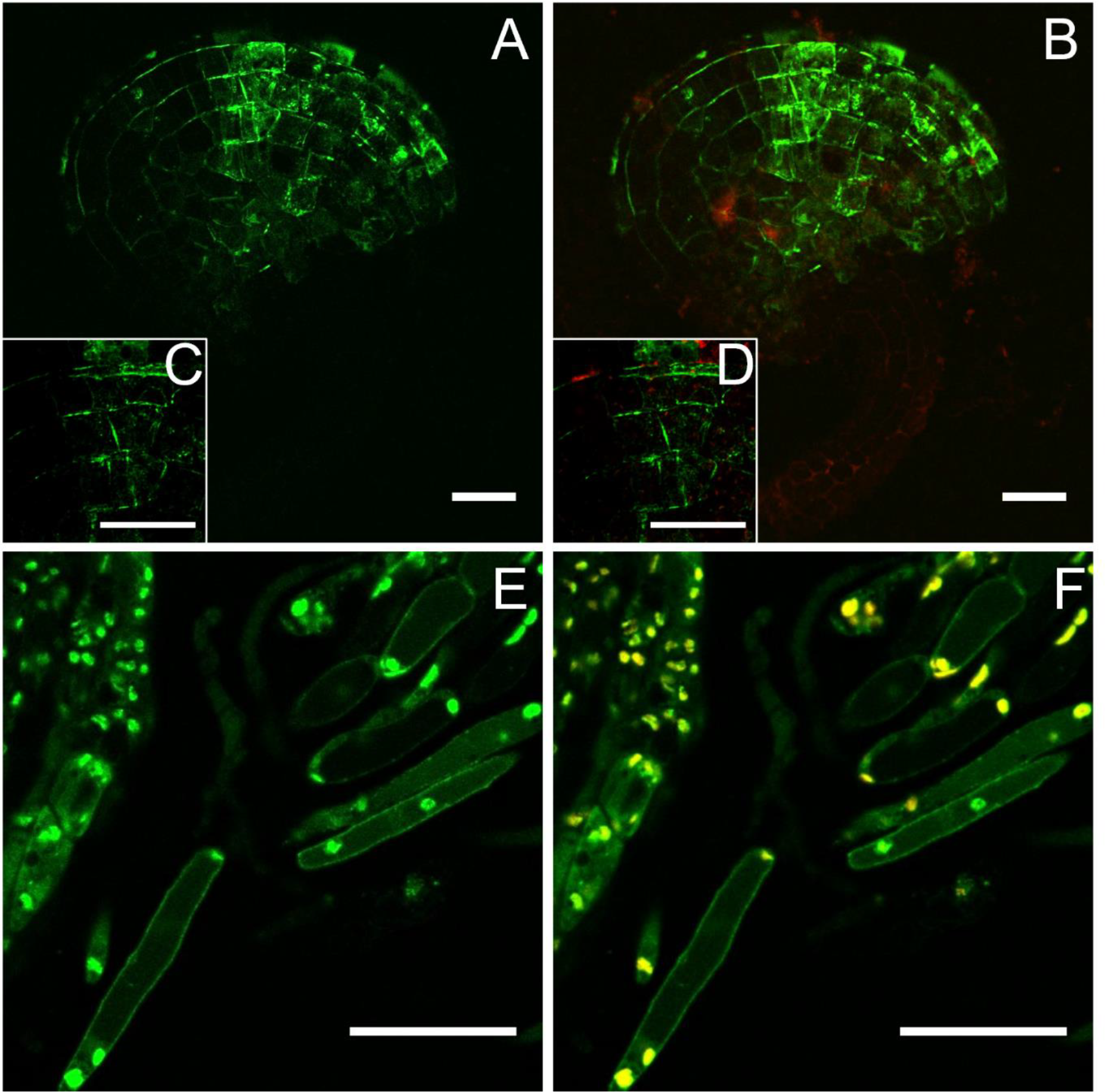
AGGER-cYFP localizes at ovule outer integument and at transmitting tissue cells **A-B**, fluorescent image showing JAGGER-cYFP localization at the ovule outer integument. **C-D**, ovule outer integument detail. E-F, fluorescent image showing JAGGER-cYFP localization at the transmitting tissue cells. A, C and E, cYFP signal. B, D and F, cYFP signal and autofluorescence merged. Scale bar 25 μm.

### *Removal of the GPI anchor attachment signal sequence in* JAGGER *negatively affects JAGGER-cYFP localization in the cell periphery*

The loss of the ω-site in JAGGER-cYFP could prevent it from receiving a GPI anchor and consequently disrupt its localization in the cell periphery. However, in ovule integuments and transmitting tissue cells in which Ser^111^ of JAGGER was deleted (JAGGER-cYFPΔω), predicted by the BIG-PI plant predictor software to have the highest chance of functioning as an ω-site in JAGGER (Eisenhaber et al. 2003), we found that the cYFP fluorescence signal was indistinguishable from that in JAGGER-cYFP (Figure 3 A – D and E – H). We hypothesized that this outcome could have been due to the redundant functions by a cryptic ω-site in JAGGER ω-region, as previously reported for LRE (Liu *et al*., 2016). The BIG-PI plant predictor software (Eisenhaber et al. 2003) identified Ser^107^ as an alternative ω-site to Ser^111^ in JAGGER, when the JAGGER protein sequence lacking Ser^111^ was used as an input at the BIG-PI plant predictor software. When both Ser^111^ and Ser^107^ were deleted, JAGGER-cYFPΔ2ω localization in the cell periphery was dramatically affected compared to wild-type JAGGER fused to cYFP (Figure 3 I – L). Concomitant with this decrease in cYFP signal in the cell periphery, there was a noticeable increase in the diffused cYFP signal in the internal regions of the cell (likely the cytoplasm) in both ovule integument cells and transmitting tissue cells of JAGGER-cYFPΔ2ω pistils (Figure 3 I - L). Further, if JAGGER protein sequence lacked both Ser^111^ and Ser^107^, the BIG-PI plant predictor software identified Ala^106^ and Ala^109^ as potential ω-sites. Removing all four predicted ω-sites, Ala^106^, Ser^107^, Ala^109^ and Ser^111^, in the JAGGER protein sequence and using it as an input in the BIG-PI plant predictor software returned no other cryptic ω-sites. The JAGGER-cYFPΔ4ω construct lacking Ala^106^, Ser^107^, Ala^109^ and Ser^111^, showed subcellular localization similar to that of JAGGER-cYFPΔ2ω, with fluorescence signal redistributed from the cell periphery to the internal regions of the cell, both in ovule integuments and transmitting tissue cells (Figure 3 M – P).

**Fig. 3.**
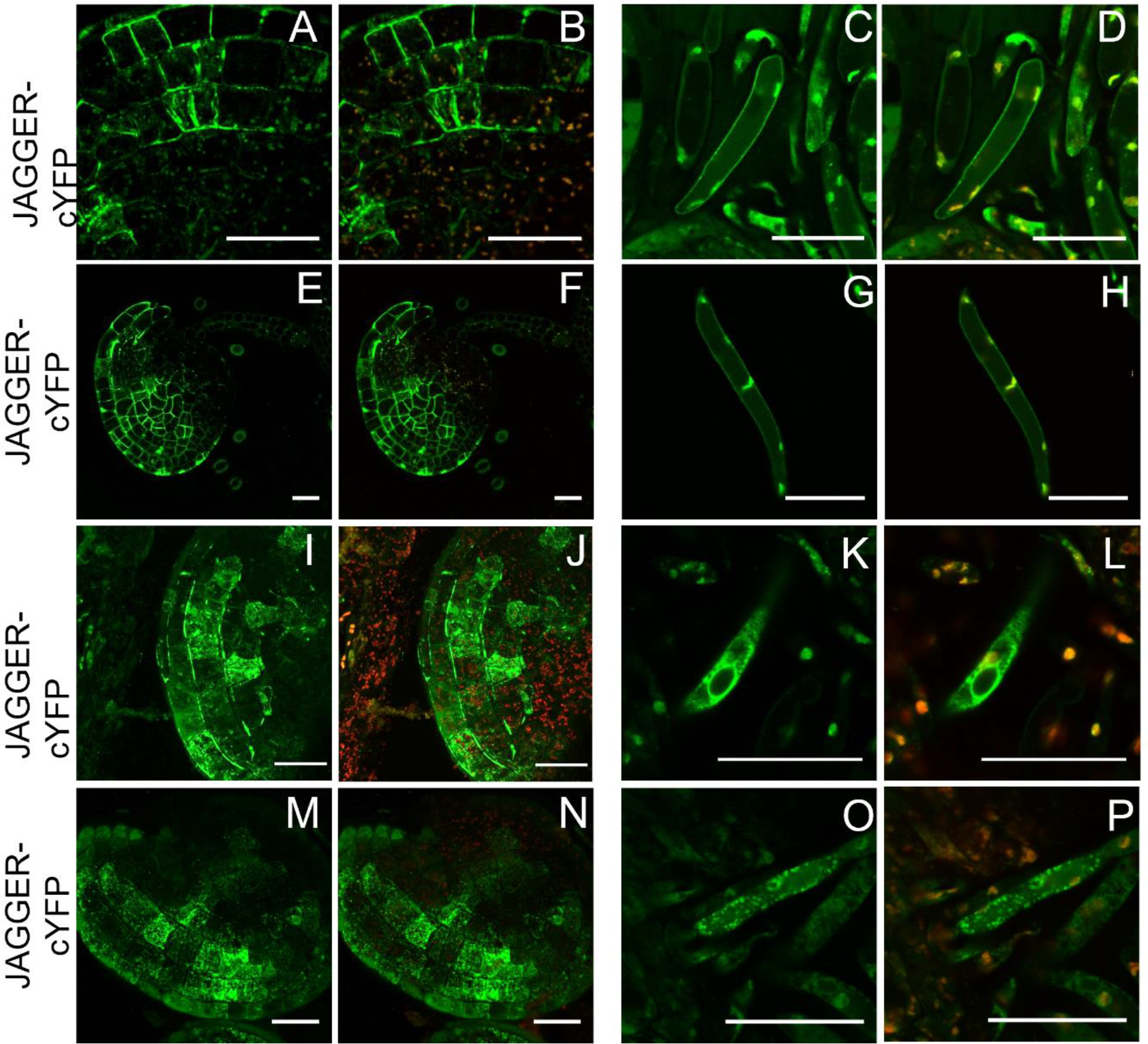
Predicted ω-site domain is not sufficient to disrupt JAGGER localization at the plasma membrane **A-D**, localization of JAGGER-cYFP in integument cells (A and B) and transmitting tract cells (C and B). **E-H**, JAGGER-cYFPΔω-site localization in ovule integuments (E and F) and transmitting tract cells (G and H). **I-L**, JAGGER-cYFPΔ2ω-site localization in ovule integuments (I and J) and transmitting tract cells (K and L). **M-P**, JAGGER-cYFPΔ4ω-site localization in ovule integuments (M and N) and transmitting tract cells (O and P). A, C, E, G, I, K, M and O, cYFP signal. B, D, F, H, J, L, N and P, cYFP signal and autofluorescence merged. Scale bar 25 μm.

The removal of putative ω-sites affected JAGGER-cYFP localization in the cell periphery; hence, we hypothesized that deletion of the GAS signal sequence (Ala^118^–Leu^134^) should also result in a non-anchored JAGGER-cYFP as predicted by (Schultz et al. 1998) and as shown for LRE (Liu et al. 2016). Contrary to these expectations, JAGGER-cYFPΔGAS^A118–L134^ showed a similar localization in the cell periphery as JAGGER-cYFP, indicating that the absence of this short C-terminal tail sequence was not sufficient to disrupt JAGGER protein sorting and localization (Figure 4 E – H). It is possible that JAGGER has a longer GAS sequence, spanning from Ser^107^ to Leu^134^, as predicted to be the case for some GAPs (Martinière et al. 2012). Transgenic plants expressing the GAS^S107–L134^ stretch of amino acid residues fused to GFP was sufficient to drive GFP to the plasma membrane in transient expression experiments performed by (Martinière et al. 2012; Bernat-Silvestre et al. 2021). Consistent with these results, we found that the localization of JAGGER-cYFPΔGAS^A109–L134^ at the cell periphery was markedly lower than JAGGER-cYFP (Figure 4 I – L) and there was a noticeable increase in the diffuse cYFP signal in the internal regions of the cells in both ovule integuments and transmitting tissue. These results were similar to the ones obtained from JAGGER-cYFPΔ2ω and JAGGER-cYFPΔ4ω localization studies and are consistent with the model in which JAGGER-cYFP localization in the cell periphery of ovule integument cells and transmitting tissue cells requires a GPI anchor.

**Fig. 4.**
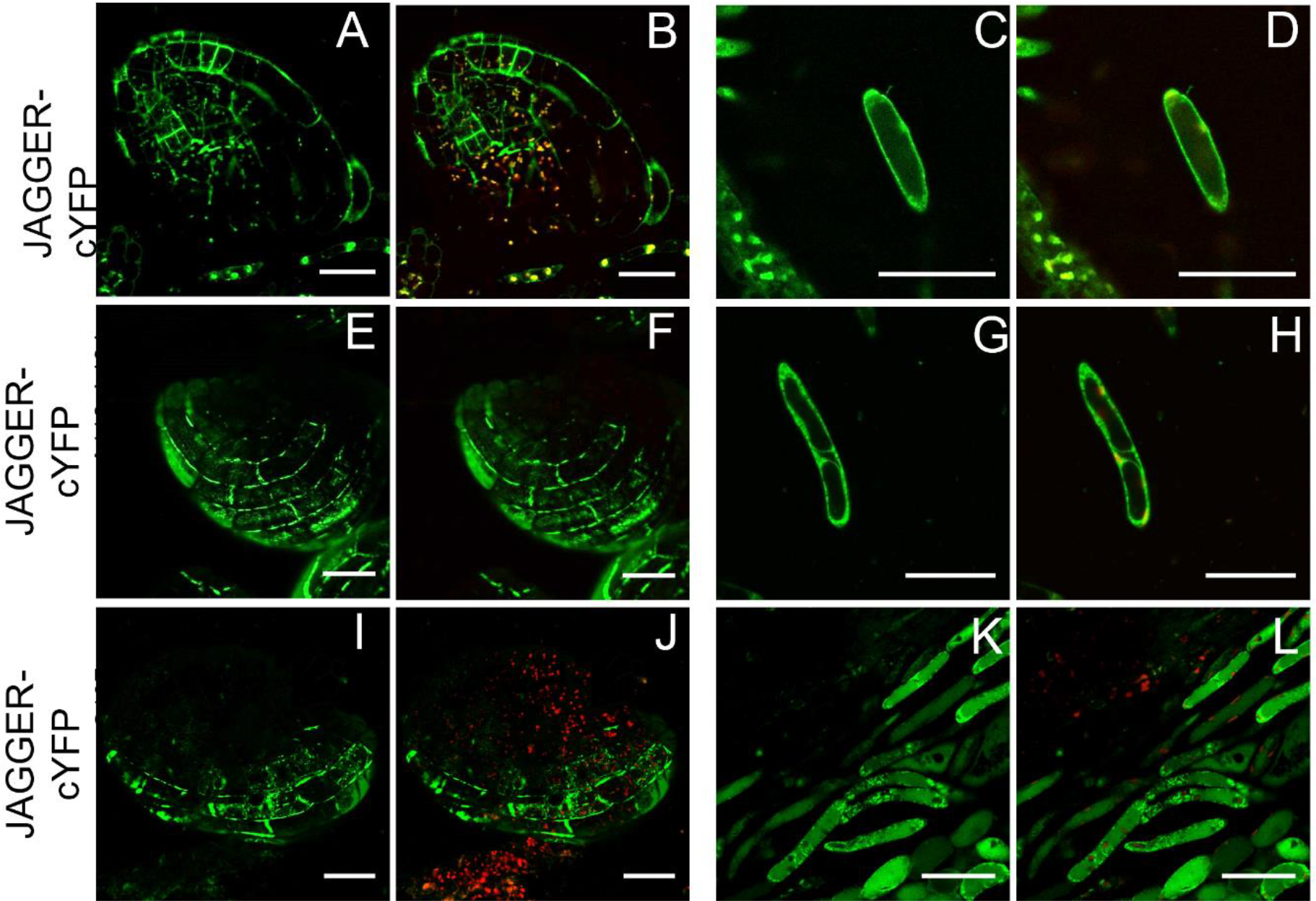
Predicted GAS^A118–L134^ domain is not sufficient to disrupt JAGGER localization at cell periphery **A-D**, localization of JAGGER-cYFP in ovule integument cells (A and B) and transmitting tract cells (C and D). **E-H**, JAGGER-cYFPΔGAS^A118–L134^ localization in ovule integument cells (E and F) and transmitting tract cells (G and H). **I-L**, JAGGER-cYFPΔGAS^S107–L134^ localization in ovule integument cells (I and J) and transmitting tract cells (K and L). A, C, E, G, I and K, cYFP signal. B, D,F, H, J and L, cYFP signal and autofluorescence merged. Scale bar 25 μm.

### The GPI anchor attachment signal sequence in JAGGER is necessary for its function in polytubey prevention

To test if indeed the GPI anchor sequence is essential for JAGGER function in polytubey block, we counted and compared the number of polytubey occurrences in wild-type hand pollinated pistils with that in *jagger-/-* pistils expressing JAGGER-cYFP and different GPI anchor mutants, namely, JAGGER-cYFPΔω, JAGGER-cYFPΔ2ω, JAGGER-cYFPΔ4ω, and JAGGER-cYFPΔ2GAS^S107–L134^ (Figure 5). Consistent with what was reported previously (Pereira et al., 2016b), polytubey was observed in 1.14% ± 1.7% of wild-type ovules (n=172). This rate of polytubey was significantly lower than that was detected in *jagger-/-* ovules (14.84% ± 1.28%; n=207). From the results shown in Figure 5, we found that in the *jagger-/-* pistils carrying constructs that did not alter JAGGER localization at the cell periphery (JAGGER-cYFP and JAGGER-cYFPΔω), there was a complete rescue of the polytubey phenotype, as the polytubey percentages in *jagger -/-* JAGGER-cYFP (2.4% ± 1.12% of ovules with polytubey occurrence, n=185) and *jagger -/-* JAGGER-cYFPΔω-site (1.87% ± 1.9% of ovules with polytubey occurrence, n=148) were similar to that observed in wild type. On the contrary, the transgenic plants that showed dramatically altered cYFP localization (*jagger -/-* plants carrying the constructs JAGGER-cYFPΔ2ω, JAGGER-cYFPΔ4ω, and the JAGGER-cYFPΔGAS^A109–L134^,) did not fully rescue the polytubey phenotype, as the polytubey percentages in these lines were 8.44% ± 3.09% (n=243), 4.69% ± 2.65% (n=229), 11.45% ± 2.47% (n=164), respectively. Importantly, when we compared the test samples analysed with the *jagger -/-* hand pollinated pistil, they were all found to have significantly different amounts of polytubey occurrences, while when we compared all the samples with wild-type hand pollinated pistils, only the *jagger -/-* pistils carrying the constructs JAGGER-cYFPΔ2ω-site, JAGGER-cYFPΔ4ω-site and the JAGGER-cYFPΔGAS^A109–L134^, presented significant differences. These results showed that the reduction in cell periphery localization of JAGGER is positively correlated with reduction in JAGGER function in preventing polytubey and point to the importance of GPI anchoring in JAGGER function.

**Fig. 5.**
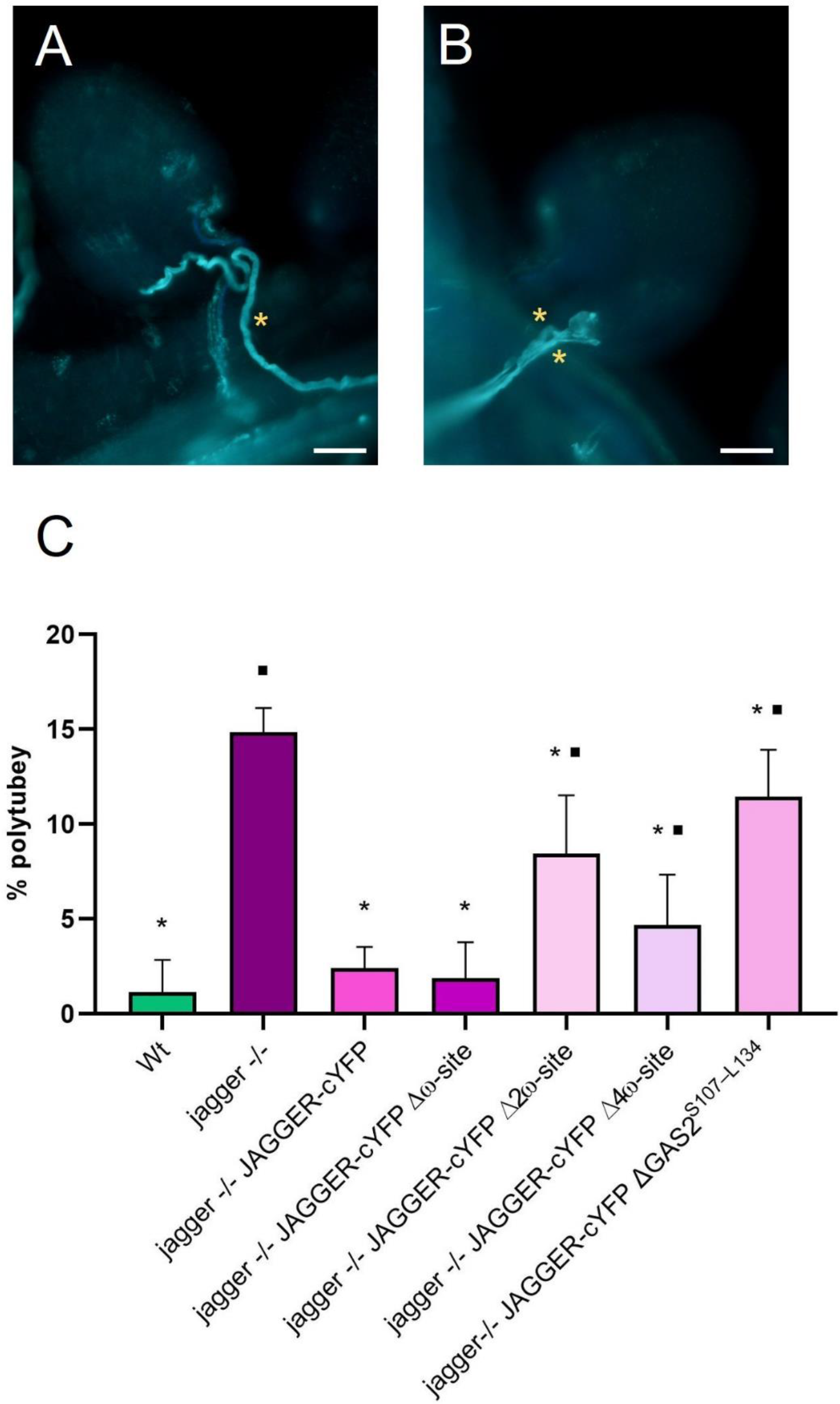
Decolorized aniline blue staining of pollen tubes growing towards the ovules of hand pollinated pistils from wild type and *jagger -/-* flowers, and the observed frequency of polytubey **A**, ovule receiving one pollen tube (yellow asterisk) from a wild type pistil. B, Ovule from a *jagger -/-* pistil receiving two pollen tubes (yellow asterisks). **C**, Frequency of polytubey observed in ovules from hand pollinated pistils stained with aniline blue in wild type (n=172), *jagger -/- (n=207), jagger-/-* JAGGER-cYFP (n=185), *jagger-/-* JAGGER-cYFPΔω-site (n=148), *jagger -/-* JAGGER-cYFPΔ2ω-site (n=243), *jagger -/-* JAGGER-cYFPΔ4ω-site (n=229) and *jagger-/-* JAGGER-cYFPΔGAS2^S107–L134^ (n=164).

## DISCUSSION

### JAGGER protein expression is consistent with the JAGGER promoter activity and its function in sporophytic tissues

JAGGER protein expression in the ovule integuments and transmitting tissue cells shown in this study is in the same region of the pistils as reported previously for JAGGER promoter fused to a reporter gene (Pereira et al., 2016b). Our protein expression data in pistils is also consistent with a sporophytic role of JAGGER in preventing polytubey (Pereira et al., 2016b). The expression results reported in this study are important because the expression of JAGGER protein in reproductive tissues has not been reported before and until now transient expression experiments performed in *Nicotiana tabacum, Nicotiana benthamiana* (Martinière et al. 2012; Zavaliev et al. 2016) and *Arabidopsis thaliana* seedlings (Bernat-Silvestre et al. 2020, 2021) used GFP-AGP4 and mCitrine-AGP4 expressed from the constitutive 35S promoter. In this study, we therefore examined the expression of the JAGGER protein in Arabidopsis pistil tissues with JAGGER-cYFP expressed from the *JAGGER* endogenous promoter.

### JAGGER is a putative GPI-anchored membrane protein in Arabidopsis pistil tissues

In the present work, we showed that JAGGER is most likely a GPI-anchored membrane protein. Until now, experimental evidence supporting JAGGER anchoring by a GPI has been reported in biochemical studies (Sherrier et al. 1999). In this study, we performed complementation tests with mutant JAGGER proteins that showed deviant subcellular localization pattern. This is important because if it is shown that the biological function of JAGGER depends on its subcellular localization, then it would be consistent with the model that its function is dependent on the presence of a GPI anchor. The confirmation that JAGGER is a GPI-AP in female tissues involved the preparation of JAGGER fusions with reporter proteins. The wild-type JAGGER protein fusions allowed us to determine the tissue and subcellular localization of JAGGER in the cells of transmitting tract and ovule integuments.

To study the importance of the GPI signal domains (present in the nascent protein) in the subcellular localization and function of JAGGER, we prepared constructs with mutations in the GPI signal domains. Subcellular localization using these proteins showed that JAGGER is a GPI anchored protein and that its subcellular localization in the cell periphery is dependent of the GPI anchor addition. The cell periphery localization of JAGGER-cYFP is also consistent with its function related to prevention of polytubey, as it involves interactions between cells in the integuments of the ovules and pollen tubes.

Another critical finding of our work is that the best predictions of the ω-site using the BIG-PI plant predictor software are not the only ω-site in this GPI-AP (Eisenhaber et al. 2003). We showed that losing the best predicted ω-site (Ser-111) in the JAGGER protein did not significantly alter its subcellular localization. In this way, we have identified another three lower score ω-sites (Ser-107, Ala-106 and Ala-109) as critical for JAGGER localization at the cell surface, suggesting that these residues may play a role in GPI anchor addition. These results are consistent with the results reported by Liu *et al*. 2016, as the deletion of a single ω-site in LORELEI-cYFP also had an indistinguishable localization from the complete LORELEI-cYFP protein, with a cryptic ω-site possibly functioning in the absence of the best predicted ω-site in LORELEI (Liu et al. 2016).

Previously, it was observed that GFP and m-Citrine fused to AGP4/JAGGER (GFP-AGP4 and mCit-AG) was exclusively localized to the cell periphery in transient expression experiments (Martinière et al. 2012; Zavaliev et al. 2016; Bernat-Silvestre et al. 2021). In the present work, it was consistently observed JAGGER-cYFP fluorescence in puncta in the cell’s cytoplasm. Still, the organelle or subcellular identity of these puncta remains unknown. Similar localization in puncta in the synergid cytoplasm was also observed with LORELEI-cYFP (Liu et al. 2016), a GPI-AP. Also, FLA4-citrine fusion revealed a similar expression in puncta of seedlings root epidermal cells. FLA4 (Fasciclin Arabinogalactan Protein 4) belongs to a subfamily of fasciclin-like AGPs, typically containing fasciclin domains and in most cases a GPI anchor (Johnson et al. 2003). These intracellular FLA4-citrine signal co-localized only partially with markers for late and recycling endosomal compartments (Xue *et al*., 2017). The authors suggest that there is a certain degree of FLA4 endocytosis, which is afterwards recycled back to the plasma membrane. This has also been proposed for AGP6 and AGP11, two pollen specific AGPs important for pollen development and pollen tube growth along the pistil tissues, based on transcriptomics and yeast two hybrid assays (Costa et al. 2013a).

Interestingly, in transient expression systems, no diffused signal in the cytoplasm has been detected for GFP-AGP4 or Venus-FLA11, another GPI-AP (Bernat-Silvestre *et al*., 2021). Although we didn’t find any study of this kind with Arabidopsis FLA4, there is one study with FLAs from *Nicotiana benthamina* which localizes FLA4 in the plasma membrane and the nucleus, in a transient expression experiment (Wu et al. 2020).

The increase in the diffuse cYFP signal in the cytoplasm of both ovule integuments and transmitting tissue cells of lines carrying JAGGER-cYFPΔ2ω and JAGGER-cYFPΔ4ω is most probably due to disruptions in its localization to the cell periphery. This raises the possibility that removal of the two predicted ω-sites led to defects in GPI anchor addition and disrupted the trafficking of JAGGER-cYFPΔ2ω to the cell periphery. If that is the case, our results may support the hypothesis that GPI attachment is required for transport of GPI-anchored proteins from the endoplasmic reticulum to the cell periphery (Mao et al. 2003). Similar disruption of localization in the plasma membrane and a concomitant diffused distribution of fluorescence in the cytoplasm happened when the entire GAS sequence in the JAGGER was removed. Most probably, the absence of the GAS region prevents its recognition and the subsequent attachment of the preassembled GPI anchor to the ω-site, supporting the importance of GPI anchor addition for JAGGER localization at the cell surface.

### GPI anchor is critical for JAGGER function in polytubey prevention

To test if losing its localization in the plasma membrane via a GPI anchor negatively affected JAGGER function, we performed several hand-pollination experiments with JAGGER-cYFP, JAGGER-cYFPΔω, JAGGER-cYFPΔ2ω, JAGGER-cYFPΔ4ω, and JAGGER-cYFPΔ2GAS constructs introduced in the *jagger-/-* background. Results from these experiments showed that only those transgenic lines in which the JAGGER localization at the cell periphery were not affected were able to rescue, at least in part, the JAGGER polytubey phenotype. These results led us to conclude that JAGGER localization in the cell periphery through a GPI anchor is critical for its function.

This result obtained for JAGGER function is strikingly different from the conclusions reported for LORELEI protein that lacked a GPI anchor, as none of the GPI anchor addition domain mutants in the LORELEI protein negatively affected LORELEI function in pollen tube reception (Liu *et al*., 2016). One possibility for this difference is that in the case of LORELEI, the nature of the filiform structure (excessive invagination of the plasma membrane and the increased presence of cell walls) could have masked the negative effects caused by the decreased concentration of mutant proteins in the cell periphery. In other words, the filiform structure in synergid cells could have accumulated enough mutant LORELEI proteins without the GPI anchor to complement the pollen tube reception phenotype. However, in case of cells that express JAGGER (transmitting tissue and ovule integuments), there are no such structural modifications to overcome the loss of JAGGER protein caused by the loss of its GPI anchor.

This result is also quite different from the one obtained for FLA4, which is implicated in many functions, including root elongation and salt stress tolerance (Xue et al. 2017). They demonstrated that FLA4 lacking the GPI anchor addition signal was partially secreted but was not membrane associated. Furthermore, this truncated version of FLA4 was still capable of complementing *sos5-1* and *sos5-2* plants phenotype under salt stress conditions. The author proposed that FLA4 may perform its function in its soluble form, rather than attached to the plasma membrane. In the case of JAGGER, as discussed in Pereira *et al*., 2016b, we suggest that it may function as a signalling molecule that interacts with receptors from the pollen tubes right before it enters the female gametophyte. In this way, the pollen tubes primed by JAGGER would act as carriers of some sort of signal for persistent synergid degeneration. And this would be in accordance with its localization at the surface of the sporophytic tissues, in this case the integuments near the micropyle entrance. Nevertheless, our results showed that mutations to GPI anchor addition domains negatively affected JAGGER function, experimentally demonstrating that GPI anchoring is key for *in vivo* functions of this AGP.

The frequency of polytubey phenotype was calculated as the number of ovules showing more than one embryo sac entering the embryo sac divided by the total number of ovules observed. Values are expressed as percentages. Error bars denote mean ± standard deviation, asterisks above bars indicate significant differences between the samples and *jagger -/-* hand pollinated pistils (P ≤ 0.05), and black squares above bars indicate significant differences between the samples and wild type hand pollinated pistils (P ≤ 0.05). One-way analysis of variance and the post hoc Tukey test were applied to detect significant differences. Statistical analysis was conducted using GraphPad Prism version 8.0.2 for Windows (www.graphpad.com). Scale bars in A and B, 20 mm.

## Author Contributions

RF and AMP performed the experiments, analysed the data, and wrote the initial draft of the manuscript. MC, DM, MM, JN and PM conducted experiments concerning construct design, plant genotyping and selection. AMP, RP, LGP and SC conceived the study, helped to analyse and interpret the data, and were involved in reviewing and writing subsequent drafts of the manuscript. All authors have read and approved the manuscript.

## Acknowledgments

This work received financial support from PT national funds (FCT/MCTES, Fundação para a Ciência e Tecnologia and Ministério da Ciência, Tecnologia e Ensino Superior) through the project UIDB/50006/2020. DM’s research was supported by an FCT PhD grant (SFRH/BD/143557/2019). SC’s research was supported by funding from an FCT SeedWheels FCT Project (POCI-01-0145-FEDER-027839). LGP research was supported by national funds through FCT - Foundation for Science and Technology within the scope of UIDB/05748/2020 and UIDP/05748/2020.

## REFERENCES

Acosta-García G, Vielle-Calzada J-P (2004) A Classical Arabinogalactan Protein Is Essential for the Initiation of Female Gametogenesis in Arabidopsis. Plant Cell 16:2614–2628. 10.1105/tpc.104.024588

Bernat-Silvestre C, De Sousa Vieira V, Sanchez-Simarro J, et al (2020) p24 Family Proteins Are Involved in Transport to the Plasma Membrane of GPI-Anchored Proteins in Plants. Plant Physiol 184:1333–1347. 10.1104/pp.20.00880

Bernat-Silvestre C, Sánchez-Simarro J, Ma Y, et al (2021) AtPGAP1 functions as a GPI inositol-deacylase required for efficient transport of GPI-anchored proteins. Plant Physiol 187:2156–2173. 10.1093/plphys/kiab384

Bundy MGR, Kosentka PZ, Willet AH, et al (2016) A Mutation in the Catalytic Subunit of the Glycosylphosphatidylinositol Transamidase Disrupts Growth, Fertility, and Stomata Formation. Plant Physiol 171:974–985. 10.1104/pp.16.00339

Capron A, Gourgues M, Neiva LS, et al (2008) Maternal Control of Male-Gamete Delivery in Arabidopsis Involves a Putative GPI-Anchored Protein Encoded by the LORELEI Gene. Plant Cell 20:3038–3049. 10.1105/tpc.108.061713

Chen X, Zaro JL, Shen W-C (2013) Fusion protein linkers: Property, design and functionality. Adv Drug Deliv Rev 65:1357–1369. 10.1016/j.addr.2012.09.039

Cheung AY, Wang H, Wu HM (1995) A floral transmitting tissue-specific glycoprotein attracts pollen tubes and stimulates their growth. Cell 82:383–393. 10.1016/0092-8674(95)90427-1

Clough SJ, Bent AF (1998) Floral dip: a simplified method for Agrobacterium -mediated transformation of Arabidopsis thaliana. The Plant Journal 16:735–743. 10.1046/j.1365-313x.1998.00343.x

Coimbra S, Almeida J, Junqueira V, et al (2007) Arabinogalactan proteins as molecular markers in Arabidopsis thaliana sexual reproduction. J Exp Bot 58:4027–4035. 10.1093/jxb/erm259

Coimbra S, Costa M, Jones B, et al (2009) Pollen grain development is compromised in Arabidopsis agp6 agp11 null mutants. J Exp Bot 60:3133–3142. 10.1093/jxb/erp148

Coimbra S, Salema R (1997) Immunolocalization of arabinogalactan proteins in Amaranthus hypochondriacus L. ovules. Protoplasma 199:75–82. 10.1007/BF02539808

Costa M, Nobre MS, Becker JD, et al (2013a) Expression-based and co-localization detection of arabinogalactan protein 6 and arabinogalactan protein 11 interactors in Arabidopsis pollen and pollen tubes. BMC Plant Biol 13:1–19. 10.1186/1471-2229-13-7/TABLES/6

Costa M, Pereira AM, Rudall PJ, Coimbra S (2013b) Immunolocalization of arabinogalactan proteins (AGPs) in reproductive structures of an early-divergent angiosperm, Trithuria (Hydatellaceae). Ann Bot 111:183–190. 10.1093/aob/mcs256

Desnoyer N, Howard G, Jong E, Palanivelu R (2020) AtPIG-S, a predicted Glycosylphosphatidylinositol Transamidase subunit, is critical for pollen tube growth in Arabidopsis. BMC Plant Biol 20:380. 10.1186/s12870-020-02587-x

Desnoyer N, Palanivelu R (2020) Bridging the GAPs in plant reproduction: a comparison of plant and animal GPI-anchored proteins. Plant Reprod 33:129–142. 10.1007/s00497-020-00395-9

Eisenhaber B, Wildpaner M, Schultz CJ, et al (2003) Glycosylphosphatidylinositol lipid anchoring of plant proteins. Sensitive prediction from sequence- and genome-wide studies for Arabidopsis and rice. Plant Physiol 133:1691–1701. 10.1104/pp.103.023580

Ellis M, Egelund J, Schultz CJ, Bacic A (2010) Arabinogalactan-proteins: Key regulators at the cell surface? Plant Physiol 153:403–419. 10.1104/pp.110.156000

Escobar-Restrepo J-M, Huck N, Kessler S, et al (2007) The FERONIA Receptor-like Kinase Mediates Male-Female Interactions During Pollen Tube Reception. Science (1979) 317:656–660. 10.1126/science.1143562

Feng H, Liu C, Fu R, et al (2019) LORELEI-LIKE GPI-ANCHORED PROTEINS 2/3 Regulate Pollen Tube Growth as Chaperones and Coreceptors for ANXUR/BUPS Receptor Kinases in Arabidopsis. Mol Plant 12:1612–1623. 10.1016/j.molp.2019.09.004

Ge Z, Cheung AY, Qu L (2019) Pollen tube integrity regulation in flowering plants: insights from molecular assemblies on the pollen tube surface. New Phytologist 222:687–693. 10.1111/nph.15645

Gjetting KSK, Ytting CK, Schulz A, Fuglsang AT (2012) Live imaging of intra- and extracellular pH in plants using pHusion, a novel genetically encoded biosensor. J Exp Bot 63:3207–3218. 10.1093/jxb/ers040

Higashiyama T, Kuroiwa H, Kawano S, Kuroiwa T (1998) Guidance in Vitro of the Pollen Tube to the Naked Embryo Sac of Torenia fournieri. Plant Cell 10:2019–2031. 10.1105/tpc.10.12.2019

Hou Y, Guo X, Cyprys P, et al (2016) Maternal ENODLs are required for pollen tube reception in Arabidopsis. Current Biology 26:2343–2350. 10.1016/j.cub.2016.06.053

Huang B-Q, Russell SD (1992) Female Germ Unit: Organization, Isolation, and Function. In: Russell SD, Dumas CBT-IR of C (eds). Academic Press, pp 233–293

Johnson KL, Jones BJ, Bacic A, Schultz CJ (2003) The Fasciclin-Like Arabinogalactan Proteins of Arabidopsis. A Multigene Family of Putative Cell Adhesion Molecules. Plant Physiol 133:1911–1925. 10.1104/pp.103.031237

Johnson MA, Harper JF, Palanivelu R (2019) A Fruitful Journey: Pollen Tube Navigation from Germination to Fertilization. Annu Rev Plant Biol 70:809–837. 10.1146/annurev-arplant-050718-100133

Karimi M, Inzé D, Depicker A (2002) GATEWAY™ vectors for Agrobacterium-mediated plant transformation. Trends Plant Sci 7:193–195. 10.1016/S1360-1385(02)02251-3

Kho YO, Baër J (1968) Observing pollen tubes by means of fluorescence. Euphytica 17:298–302. 10.1007/BF00021224

Lamport DTA, Tan L, Kieliszewski MJ (2021) A Molecular Pinball Machine of the Plasma Membrane Regulates Plant Growth—A New Paradigm. Cells 10:1935. 10.3390/cells10081935

Lara-Mondragón CM, MacAlister CA (2021) Arabinogalactan glycoprotein dynamics during the progamic phase in the tomato pistil. Plant Reprod 34:131–148. 10.1007/s00497-021-00408-1

Li H-J, Meng J-G, Yang W-C (2018) Multilayered signaling pathways for pollen tube growth and guidance. Plant Reprod 31:31–41. 10.1007/s00497-018-0324-7

Li J, Yu M, Geng L-L, Zhao J (2010) The fasciclin-like arabinogalactan protein gene, FLA3, is involved in microspore development of Arabidopsis. Plant J 64:482–497. 10.1111/j.1365-313X.2010.04344.x

Li S, Ge F-R, Xu M, et al (2013) Arabidopsis COBRA-LIKE 10, a GPI-anchored protein, mediates directional growth of pollen tubes. The Plant Journal 74:486–497. 10.1111/tpj.12139

Liu X, Castro C, Wang Y, et al (2016) The Role of LORELEI in Pollen Tube Reception at the Interface of the Synergid Cell and Pollen Tube Requires the Modified Eight-Cysteine Motif and the Receptor-Like Kinase FERONIA. Plant Cell 28:1035 LP –1052. 10.1105/tpc.15.00703

Lopes AL, Costa ML, Sobral R, et al (2016) Arabinogalactan proteins and pectin distribution during female gametogenesis in Quercus suber L. Ann Bot 117:949–961. 10.1093/aob/mcw019

Lopes AL, Moreira D, Ferreira MJ, et al (2019) Insights into secrets along the pollen tube pathway in need to be discovered. J Exp Bot 70:2979–2992. 10.1093/jxb/erz087

Majewska-Sawka A, Nothnagel EA (2000) The Multiple Roles of Arabinogalactan Proteins in Plant Development1. Plant Physiol 122:3–10. 10.1104/pp.122.1.3

Mao Y, Zhang Z, Wong B (2003) Use of green fluorescent protein fusions to analyse the N- and C-terminal signal peptides of GPI-anchored cell wall proteins in Candida albicans. Mol Microbiol 50:1617–1628. 10.1046/j.1365-2958.2003.03794.x

Martinière A, Lavagi I, Nageswaran G, et al (2012) Cell wall constrains lateral diffusion of plant plasma-membrane proteins. Proceedings of the National Academy of Sciences 201202040. 10.1073/pnas.1202040109

Mizukami AG, Inatsugi R, Jiao J, et al (2016) The AMOR arabinogalactan sugar chain induces pollen-tube competency to respond to ovular guidance. Current Biology 26:1091–1097. 10.1016/j.cub.2016.02.040

Moreira D, Kaur D, Pereira AM, et al (2023) Type II arabinogalactans initiated by hydroxyproline-O-galactosyltransferases play important roles in pollen–pistil interactions. The Plant Journal 114:371–389. 10.1111/tpj.16141

Oxley D, Bacic A (1999) Structure of the glycosylphosphatidylinositol anchor of an arabinogalactan protein from Pyrus communis suspension-cultured cells. Proc Natl Acad Sci U S A 96:14246–14251. 10.1073/pnas.96.25.14246

Pereira AM, Coimbra S (2019) Advances in plant reproduction: from gametes to seeds. J Exp Bot 70:2933–2936. 10.1093/jxb/erz227

Pereira AM, Lopes AL, Coimbra S (2016a) Arabinogalactan Proteins as Interactors along the Crosstalk between the Pollen Tube and the Female Tissues. Front Plant Sci 7:1895. 10.3389/fpls.2016.01895

Pereira AM, Nobre MS, Pinto SC, et al (2016b) “Love is strong, and you’re so sweet”: JAGGER is essential for persistent synergid degeneration and polytubey block in Arabidopsis thaliana. Mol Plant 9:601–614. 10.1016/j.molp.2016.01.002

Qin Y, Chen D, Zhao J (2007) Localization of arabinogalactan proteins in anther, pollen, and pollen tube of Nicotiana tabacum L. Protoplasma 231:43–53. 10.1007/s00709-007-0245-z

Russell SD (1992) Double Fertilization. In: Russell SD, Dumas CBT-IR of C (eds). Academic Press, pp 357–388

Schultz C, Gilson P, Oxley D, et al (1998) GPI-anchors on arabinogalactan-proteins: Implications for signalling in plants. Trends Plant Sci 3:426–431. 10.1016/S1360-1385(98)01328-4

Seifert GJ, Roberts K (2007) The Biology of Arabinogalactan Proteins. Annu Rev Plant Biol 58:137–161. 10.1146/annurev.arplant.58.032806.103801

Sherrier DJ, Prime TA, Dupree P (1999) Glycosylphosphatidylinositol-anchored cell-surface proteins from Arabidopsis. Electrophoresis 20:2027–2035. 10.1002/(SICI)1522-2683(19990701)20:10<2027::AID-ELPS2027>3.0.CO;2-A

Shimizu KK, Okada K (2000) Attractive and repulsive interactions between female and male gametophytes in Arabidopsis pollen tube guidance. Development 127:4511–4518. 10.1242/dev.127.20.4511

Simpson C, Thomas C, Findlay K, et al (2009) An Arabidopsis GPI-anchor plasmodesmal neck protein with callose binding activity and potential to regulate cell-to-cell trafficking. Plant Cell 21:581–594. 10.1105/tpc.108.060145

Smyth DR, Bowman JL, Meyerowitz EM (1990) Early flower development in Arabidopsis. Plant Cell 2 (8): 755–767. 10.1105/tpc.2.8.755

Tian G-W, Mohanty A, Chary SN, et al (2004) High-throughput fluorescent tagging of full-length Arabidopsis gene products in planta. Plant Physiol 135(1): 25–438

Wu H, Wang H, Cheung AY (1995) A pollen tube growth stimulatory glycoprotein is deglycosylated by pollen tubes and displays a glycosylation gradient in the flower. Cell 82:395–403. 10.1016/0092-8674(95)90428-X

Wu X, Lai Y, Lv L, et al (2020) Fasciclin-like arabinogalactan gene family in Nicotiana benthamiana: genome-wide identification, classification and expression in response to pathogens. BMC Plant Biol 20:305. 10.1186/s12870-020-02501-5

Xue H, Veit C, Abas L, et al (2017) Arabidopsis thaliana FLA4 functions as a glycan-stabilized soluble factor via its carboxy-proximal Fasciclin 1 domain. The Plant Journal 91:613–630. 10.1111/tpj.13591

Yadegari R, Drews GN (2004) Female Gametophyte Development. Plant Cell 16:S133 LP–S141. 10.1105/tpc.018192

Yeats TH, Bacic A, Johnson KL (2018) Plant glycosylphosphatidylinositol anchored proteins at the plasma membrane-cell wall nexus. J Integr Plant Biol 60:649–669. 10.1111/jipb.12659

Zavaliev R, Dong X, Epel BL (2016) Glycosylphosphatidylinositol (GPI) modification serves as a primary plasmodesmal targeting signal. Plant Physiol pp.01026.2016. 10.1104/pp.16.01026

