## Supplemental Table 1 for "JAGGER localization and function is dependent on GPI anchor addition"

Supporting Table 1 - List of primers used in this work

| Name | Sequence | Description |
|---|---|---|
| *ProJAGGER:JAGGER-cYFP* | | |
| JAGGER::cYFP - 1 | GGCGGCCGCACTAGTCTGGTAATATTTGATAAAAGCGG | Anneals in JAGGER promoter; includes a 15 bp overlap with modified pH7WG vector (red). Fw |
| JAGGER::cYFP - 2 | GGCCTGGGGCAGGGGATGCATCTGA | Anneals in JAGGER ω-11 region; includes a 15 bp overlap with primer JAGGER::cYFP - 3 (red). Rv |
| JAGGER::cYFP - 3 | CCCCTGCCCCAGGCCGGCCTGGAGGTGGA | Anneals in $Gly^9$ linker::cYFP construct (plasmid E1403); includes a 15 bp overlap with primer JAGGER::cYFP - 2 (red). Fw |
| JAGGER::cYFP - 4 | TGGAGAAGCGTCAGATGGACCAGGTGCCTTGTACAGCTCGTCCATG | Anneals in $Gly^9$ linker::cYFP construct (plasmid E1403); includes a revised ω-11 overhang (red) which last 15 bp overlap with primer JAGGER::cYFP - 5. Rv |
| JAGGER::cYFP - 5 | TCTGACGCTTCTCCAGCACCTAGCGCCGCATTCTCC | Anneals in JAGGER coding sequence; includes a revised ω-11 overhang (red) which first 15 bp overlap with primer JAGGER::cYFP - 4. Fw |
| JAGGER::cYFP - 6 | CTGGGTCGGCGCGCCATGTTTAAGTGAAAGAGTAATGAAATTGAT | Anneals in JAGGER 3'UTR; includes a 15 bp overlap with modified pH7WG vector (red). Rv |
| *ProJAGGER:JAGGER-cYFP Δω* | | |
| Δω-site - 1 | AGAATGCGGCAGGTGCTGGAGAAGCGTC | Pairs with JAGGER::cYFP - 1; Anneals in JAGGER ω-11 region and incorporates deletion of the highest predicted ω-site (S111). Rv |
| Δω-site - 2 | AGCACCTGCCGCATTCTCCAACAAGG | Pairs with JAGGER::cYFP - 6; Anneals in JAGGER ω-11 region and incorporates deletion of the highest predicted ω-site (S111). Fw |
| *ProJAGGER:JAGGER-cYFP Δ2ω* | | |
| Δ2ω-site - 1 | GCAGGTGCTGGAGCGTCAGATGGACCAGG | Pairs with JAGGER::cYFP - 1; Anneals in JAGGER ω-11 region and incorporates deletion of the two highest predicted ω-site (S111; S107). Rv |
| Δ2ω-site - 2 | CGCTCCAGCACCTGCCGCATTCTCCAACAAGGC | Pairs with JAGGER::cYFP - 6; Anneals in JAGGER ω-11 region and incorporates deletion of the two highest predicted ω-site (S111;S107). Fw |
| *ProJAGGER:JAGGER-cYFP ΔGAS* | | |
| ΔGAS - 1 | TAAAAAAAGTTCTCAAGCCTTGTTGGAGAATGCGGCGCT | Pairs with JAGGER::cYFP - 1; Anneals in JAGGER CDS+3'UTR and incorporates deletion of the hydrophobic GAS domain (A118-L134). Rv |
| ΔGAS - 2 | GCTTGAGAACTTTTTTTATATAATTTTTTTTTTTATCCCTCAAATT | Pairs with JAGGER::cYFP - 6; Anneals in JAGGER CDS+3'UTR and incorporates deletion of the hydrophobic GAS domain (A118-L134). Fw |
| *ProJAGGER:JAGGER-cYFP Δ4ω* | | |
| Δ4ω-site - 1 | CGGCAGGTGGGTCAGATGGACCAGGTGCC | Pairs with JAGGER::cYFP - 1; Anneals in JAGGER ω-11 region and incorporates deletion of the 4 highest predicted ω-site (S111; S107; A109; A106). Rv |
| Δ4ω-site - 2 | CTGACCCACCTGCCGCATTCTCCAACAAGGC | Pairs with JAGGER::cYFP - 6; Anneals in JAGGER ω-11 region and incorporates deletion of the two highest predicted ω-site (S111;S107; A109; A106). Fw |
| *jagger::ProJAGGER:JAGGER-cYFP and deletions genotyping* | | |
| LP-GK-134A10 | TGTCTCCCCACATTTGCCAT | Pairs with RP-GK-134A10; Anneals in JAGGER promoter. Fw |
| RP-GK-134A10 | ACAACCATATGAAGCCCTTCC | Pairs with LP-GK-134A10; Anneals in JAGGER 3'UTR. Rv |
| cYFP_Fw | ACGACGGCAACTACAAGACC | Anneals in cYFP; used to sequence the 3' end of *JAGGER-CYFP* constructs. Fw |
